# Localised stabilisation of a bistable switch for minimal-kernel phenotype control in a Boolean breast-cancer network

**DOI:** 10.64898/2026.06.18.733086

**Authors:** Aamer Iqbal Bhatti

## Abstract

We pose breast-cancer therapy as a control problem on a Boolean network of signalling pathways. The goal is a minimal, druggable intervention that drives the network to apoptosis as a genuine fixed point of the free dynamics, with no node held on by force. On a 141-node network, adapted substantially from Taoma et al. [22] with changes to its feedback wiring, a capped, no-forcing whole-network controller shows that the cell-fate phenotypes are individually controllable but jointly near-exclusive. They share a dominant strongly-connected core, 16 of 22 pathways, so feasibility does not imply joint reachability. We therefore reduce the apoptosis core to a nine-node bistable switch on the AKT1 and TP53 axis. We embed biological timescale separation as a multirate Boolean switch, fast, mid, and slow on a 1:5:25 schedule, with the slow variables sampled and held. We then relax global stabilisation, which is intractable on the full network, to localised stabilisation on the switch. A minimal kernel of about two to three nodes, concentrated on the PI3K/AKT/mTOR axis, drives the switch to its apoptotic attractor. It commits the full 141-node network to a caspase-cascade death fixed point in 15 to 17 percent of patients across three cohorts (TCGA, METABRIC, I-SPY2). It holds that state as durably as a near whole-network controller of about thirty nodes, an order-of-magnitude reduction at equal durability. The model is a structural drug-target nominator, not a response predictor.

## 1 Introduction

Targeted agents in breast cancer often shrink a tumour, then lose durable control of it. The drug blocks a signalling hub. The pathway relieves its own brakes. The hub reactivates. The most recent approved agent in this class delays relapse rather than preventing it. In CAPItello-291, progression-free survival rose from 3.1 to 7.3 months, yet the disease still advanced [26]. The obstacle is not reaching an apoptotic state. It is making that state durable. Durability is the gap under current treatment. One way a model manufactures a durable death state is to pin nodes at fixed values. That is a theoretical device, not a therapy. No drug holds a protein clamped in a patient. So a durable state that depends on sustained pinning does not survive contact with the clinic. The durability that does survive comes from the dynamics. Drive the system into the apoptotic basin of attraction, the set of states from which the free dynamics settle to death on their own, and it stays there once the drive is removed. We therefore treat therapy as a control problem. On a Boolean network of the relevant signalling, we seek a minimal, druggable intervention. It must drive the network into that basin, so the apoptotic state is a genuine fixed point of the free dynamics, not one held on by force.

Boolean networks suit this question. They capture the logic of regulation without kinetic constants that are rarely measurable [12, 13]. Their predictive value for real biology is established [14]. Every trajectory settles into an attractor, and each attractor maps to a cell fate such as proliferation or apoptosis. Logic-based modelling of signalling is now systematised, with mature tools for large models [15, 16, 17, 18]. In cancer, such models reproduce the hallmarks and test interventions [19, 20, 21, 22, 23]. The therapeutic question becomes one of control. Which nodes should be perturbed, and how, to steer the network from a pathological attractor to a desired one?

Two traditions bear on that question. Our approach is defined by where it leaves each. The first is structural attractor control. Stable-motif [3, 4, 5] and feedback-vertex-set [24] methods certify from topology that a target attractor is controllable in principle. The second is algebraic control by the semi-tensor product [6, 35]. It renders a Boolean control network as a linear map over a lifted state space. Controllability, stabilisation, and their time-delayed extensions all follow [34, 36, 37]. This power has one main cost. The lifted space is exponential in the number of nodes, and a delay multiplies it further. For a network the size of a real signalling model the semi-tensor-product form is too large to build, let alone to design a controller on. Global stabilisation by this route is then a theoretical object more than a usable one. A separate strand fits a generic model to patient omics and simulates drug effects [7, 8]. These are all powerful. But none of them yields a controller under a realistic budget. The simulators annotate rather than design. The structural methods certify that a target is controllable in principle, but not with a bounded, druggable set of interventions. The algebraic methods, where they stay tractable, deliver global stabilisation, which pins far more of the network than any therapy could reach. What is missing is control design under a cap, a small, non-accumulating, druggable intervention on real patient states. This paper is that design.

Two obstacles stand between feasibility and a deliverable controller. The first is joint reachability. Individual phenotypes can each be driven to a fixed point. Reaching them together under a capped budget can still fail. The cause is a dominant strongly-connected core shared across pathways. The second obstacle is time. The standard Boolean update advances every node on one clock. The apoptosis core does not work that way. Fast survival signalling and slow transcriptional feedback decide fate together. Forced onto one clock, the core oscillates, and no durable death state is reachable. Different timescales must be built in. This is an established device in asynchronous and priority-class Boolean modelling [38, 39]. Here it is a precondition for control of the core, not an embellishment.

We address both obstacles on one network. First, a capped, no-forcing whole-network controller is run on three cohorts. It tests reachability, not feasibility. The phenotypes are individually reachable but jointly near-exclusive. A dominant strongly-connected core, 16 of 22 pathways, obstructs joint control. This result forces the retreat to the core. It is not a convenience. Second, we reduce the apoptosis core to a nine-node bistable switch on the AKT1 and TP53 axis.

The control-design idea is a relaxation. Global stabilisation asks every node of the full network to converge to the target [2]. That is intractable here, and it is unnecessary. We use local stabilisation instead [1]. A target set is locally stabilised when trajectories from its neighbourhood converge to it under the free dynamics. This holds when the coupling into the target block is not significant. Whether that condition holds depends on how the network is partitioned into local subsystems. In our earlier modular design, each pathway module was one local subsystem [1], and the coupling between pathway modules was not always weak. Here we partition by timescale instead. Each time-delayed block, fast, mid, or slow, is one local subsystem. Two things then reduce the coupling. The delay structure separates the processes in time. And the unit of design is now the timescale block, not the pathway module. Between these blocks the coupling is weak. The local-stabilisation condition holds block by block. We can therefore design a controller for each block on its own. This is what makes a semi-tensor-product design usable here. A global design must lift the whole 141-node state space, and that algebraic dimension grows exponentially, worsened further by delay [37]. Block-wise design avoids that dimension explosion. It confines the semi-tensor product to a small switch, where it stays tractable. The result relaxes the weak-coupling pathway decomposition of our earlier work [1]. Sequential stabilisation across all pathways becomes a single localised action on a bistable core.

The kernel this yields is the answer to the durability problem. It is minimal and druggable, about two to three nodes, concentrated on the PI3K/AKT/mTOR axis. Its job is not to hold the death state in place. Its job is to drive the system across the barrier into the apoptotic basin. The free dynamics then keep it there, with no sustained forcing. A capped kernel found by our search is enough to do this, and the state it reaches is held as durably as a near whole-network controller of about thirty nodes. These are not speculative targets. The PI3K, AKT, and mTOR axis is already among the most actively drugged in breast cancer, so the kernel the design returns is immediately actionable rather than hypothetical.

The network itself is not a contribution. It is adapted substantially from the breast-cancer Boolean model of Taoma and colleagues [22]. We adapt that model. Two feedback loops are closed: the p53 loop, which carries a DNA-damage input, and the AKT1 and FOXO3a loop, closed by PHLPP. The intrinsic caspase cascade is then added, so that apoptotic commitment becomes irreversible. Methods gives the detail, and the appendix lists the full model. We treat this as model construction, not as a finding.

This paper makes one methodological contribution, with four supporting results. The contribution is a control-design principle. On a strongly coupled Boolean cancer network, global stabilisation is infeasible. Localised stabilisation of an extracted bistable switch is enough. It steers the whole network’s phenotype with a minimal, druggable kernel. The supporting results are four. First, cell-fate phenotypes are individually controllable but jointly near-exclusive, obstructed by a dominant strongly-connected core. Second, a multirate Boolean embedding, with the slow variables sampled and held, makes the switch’s apoptotic state reachable and stable, where a single-clock model only oscillates. Third, a capped switch-designed kernel of two to three nodes drives the network into the apoptotic basin, which then holds as durably as near whole-network control of thirty. Fourth, resistance is a hysteresis property. Its control target is the AKT1 and FOXO3a coupling state, and survival-axis inhibition flips exactly the coupled fraction. The model is a structural drug-target nominator, not a response predictor. We state that scope plainly and return to it in the Discussion. The paper is organised as follows. Methods covers the cohorts and binarisation, the Taoma-derived 141-node network and its adaptations, the capped whole-network controller, the delayed multirate switch, and the localised-stabilisation kernel design. Results reports the joint-reachability obstruction (Section 3.1), the switch and its patient routing (Section 3.2), the localised-stabilisation controller and the apoptosis it secures on the full network (Section 3.4), and the minimality of that controller (Section 3.5). The Discussion places localised stabilisation against stable-motif and semi-tensor-product control theory, and states the limits of the claim. Every number, figure, and table regenerates from the public repository with one command.

## 2 Methods

### 2.1 Cohorts and binarisation

Patient states come from primary-tumour mRNA in three public cohorts. TCGA-BRCA [10] has 1082 patients on RNA-seq. METABRIC [11] has 1980 on microarray. I-SPY2 has 988 on microarray, from GEO series GSE194040. The three span different platforms and different patient populations. A finding that holds across all three is therefore unlikely to be an artefact of one assay or one centre. TCGA and METABRIC are large reference cohorts on unrelated technologies. I-SPY2 is a neoadjuvant trial cohort that also carries treatment annotation.

Each gene is binarised within its own cohort. The activity score is the robust *z*-score, (*x* − median)*/*(1.4826 · MAD), taken over all patients in the cohort. The node is set ON when the score exceeds zero. The robust estimator is the default. It uses the median and the median absolute deviation, not the mean and the standard deviation, so a few extreme expression values cannot drag the threshold. Such values are common in skewed RNA-sequencing data. Classic *z*-score and hard-median splits are available as alternatives. Some nodes stand for a signalling activity rather than one gene. For these a curated multi-gene panel is reduced to one bit by majority vote: the node is ON when at least half of its panel genes are ON. The panel memberships are part of the model source. The resulting Boolean vector, restricted to the network nodes, is the patient’s initial state.

Three features of this encoding should be kept in view. The threshold is the cohort median, so each gene is ON in about half of the cohort by construction. This is a deliberate coarse-graining, suited to a qualitative Boolean model rather than to absolute expression levels. Binarisation is within each cohort, so a state is defined relative to that cohort, and agreement across cohorts is concordance of behaviour, not of absolute activity. And mRNA stands in for signalling activity, which for kinases such as AKT is set after translation, so the initial state is a transcriptional proxy, not a direct activity measurement.

### 2.2 The network: an inherited and extended model

The network is not a contribution of this paper, and we describe it as construction. It is adapted substantially from the breast-cancer Boolean model of Taoma and colleagues [22]. Of the 135 nodes in our base model, 105, or 78 percent, correspond to nodes in that model. Once Taoma’s merged complexes and isoforms are re-expressed at individual-gene resolution, for example AKT1, 2, and 3 as one node there and separate nodes here, the update rules coincide for the majority of the shared nodes. The apoptosis and proliferation readouts, CASP3 and E2F1, follow Taoma’s convention.

We extend that base from 117 to 135 nodes across 24 pathway modules. The additions are a DNA-damage arm (ATM, ATR, BRCA1, BRCA2, CHEK1, CHEK2), immune-checkpoint nodes, and NF-kB nodes. The immune-checkpoint and NF-kB nodes are carried for completeness of the inherited network; the results of this paper exercise the apoptosis core, not those arms. The complete rule set of the full model is listed in Appendix A. The work also builds on the author’s earlier BBCN118 framework [1].

We adapt the model in two respects, each a piece of established apoptosis biology. Two feedback loops are closed, and one commitment mechanism is added. We describe them as biology. Separately, we note that together they are what let a durable apoptotic fixed point exist, which Section 3.4 then tests, rather than a target we tuned the model toward.

The first is the p53 and MDM2 loop [40, 41]. We represent the negative feedback together with a DNA-damage input. MDM2 is driven by TP53 and AKT1 and relieved by the CDKN2A and E2F1 damage arm. TP53 is raised by ATM and ATR. This closes the p53 response to damage. It adds no node.

The second is the AKT1 and FOXO3a loop [32, 42]. We close it with PHLPP, the phosphatase that turns AKT1 back off once FOXO3a is nuclear. PHLPP tracks FOXO3, and PHLPP represses AKT1. This closes the loop. It adds one node, PHLPP, taking the model to 136 nodes.

The third change is different in kind. The baseline has no irreversible commitment step, so an apoptotic excursion can reverse. We add the intrinsic caspase cascade, whose bistable, self-sustaining behaviour is well characterised [27, 28, 29]. p53 induces PUMA and NOXA [43, 44], which repress the anti-apoptotic guardians. BAX and BAK release SMAC, which neutralises XIAP [45]. An executioner loop then closes: CASP6 activates CASP8, CASP8 activates CASP3, and CASP3 feeds back to CASP6. Once an executioner fires it self-sustains, the molecular point of no return. The change adds five nodes, PUMA, NOXA, SMAC, CASP6, and CASP7, taking the model to its full size of 141 nodes. It is an addition, not the repair of a collapsed loop, though the executioner loop it introduces is itself a feedback structure.

We treat these adaptations as model construction, not as findings of this work. Their role is to let a durable apoptotic fixed point exist, which Section 3.4 then shows the localised controller can reach.

A note on model construction. This is a Boolean model, so it has no fitted parameters or thresholds to tune. Its content is structural: which nodes, and which logic rules. The network structure was fixed on biological grounds, and the cohort analyses were run against the finished model rather than used to shape it. The additions above are corrections against established apoptosis biology, each independently cited, not adjustments made to obtain a result.

### 2.3 The whole-network controller

The whole-network controller is the instrument that reveals the obstruction. Its job in this paper is to test reachability on the full network under a realistic budget, and so to show why the retreat to the switch is necessary. We describe it here. The controller is specified exactly by the released code, in `setup_a/` of the companion repository.

The controller has three tiers, each on its own unit of time and decision. The top tier sequences the target phenotypes in clinical order: resistance off, then apoptosis on, then proliferation off. The middle tier acts on a clinical cycle of twenty-one days, and its unit is pathways. From a candidate set it ranks pathways, upstream first, and passes one or two down. The bottom tier acts on a single network step, taken as one day, and its unit is kernels. For each selected pathway it designs the minimal stabilising kernel and administers it. A cycle is one design step followed by free relaxation. The kernel is fixed at the start of the cycle, and the network then evolves freely under it. The horizon is eight cycles. The day and cycle lengths are an illustrative clinical cadence; the obstruction does not depend on them.

The intervention is capped and does not accumulate. During induction up to two pathways act at once, which pins four to five nodes. During maintenance one pathway acts, which pins two or three. The kernel is re-decided at each cycle boundary and never exceeds the cap. Prior-cycle pins are not retained. The bus state carries forward, but the intervention itself does not. This keeps the modelled therapy within a plausible drug-combination budget, and it stops the controller from quietly clamping the whole network. The obstruction we report is therefore a statement about reachability under a deliverable budget. A larger budget would pin more of the network; the point of the cap is that a therapy cannot.

The kernel at each step is found by the same algebraic forward-stabilisation test [2] used in Section 2.5, applied here at the granularity of the pathway module, with the per-module kernels composed in sequence, in the upstream-first order of the middle tier. A different order could compose differently, so the result is evidence of difficulty under a deliverable, ordered budget, not a proof of impossibility. This is the pathway-module partition. It is the partition whose weak-coupling premise Section 2.5 shows to fail on the apoptosis core, and the timescale partition replaces it. Run this controller on the three cohorts and the phenotypes are individually reachable but jointly near-exclusive. That obstruction, and the strongly-connected core behind it, are reported in Section 3.1.

### 2.4 The apoptosis core as a bistable switch

The apoptosis decision is carried by a small, well-studied antagonism. AKT1 drives survival. TP53 drives death. Each represses the other, AKT1 through MDM2 and TP53 through PHLPP and FOXO3a [27, 28, 32]. We take the nine nodes that carry this antagonism as a reduced switch. The reduction is standard and is not a contribution of this paper. The contribution is the control design on it (Section 2.5). The nodes and their timescale classes are given in Table 1: five fast (phosphorylation), one mid (protein turnover), and three slow (transcription). The rest of the network enters only through a held, patient-derived input snapshot *I*, read once from the binarised state. The full Boolean rules are listed in Appendix A.

**Table 1.** The nine switch nodes, their timescale class, update rate, and role. The three classes advance in the ratio 1:5:25, and the slow nodes enter the fast rules at a delay.

| Node | Class | Rate | Role |
| --- | --- | --- | --- |
| AKT1 | fast | 1 | survival hub |
| PIK3CA | fast | 1 | PI3K catalytic input |
| PTEN | fast | 1 | PI3K/AKT brake |
| PDPK1 | fast | 1 | AKT activator |
| MTOR | fast | 1 | growth signalling |
| MDM2 | mid | 5 | p53 repressor |
| TP53 | slow | 25 | apoptosis hub |
| PHLPP | slow | 25 | AKT phosphatase, closes the AKT and FOXO3a loop |
| FOXO3 | slow | 25 | pro-apoptotic transcription factor |

Time enters through the update schedule, not the rules. The three classes advance at rates 1:5:25. These are not fitted values. They stand for the order-of-magnitude separation between the processes: phosphorylation acts in seconds to minutes, protein turnover in minutes to hours, and transcription in hours [38, 39]. Only the separation of scales matters for the switch, not the exact integers, and a finer calibration is left to future work. A fast rule that reads a slow node reads its value from the last slow update, a sample and hold. Writing the fast state as *x*_*f*_ and the slow state as *x*_*s*_,

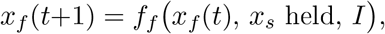

with the slow block held constant over its own update interval. Over a fast interval the slow block is a fixed input, and this is the structure that Section 2.5 uses to make the blocks weakly coupled.

The switch is bistable, and we verify this directly rather than assume it. Enumerating the 2^9^ states, the free dynamics settle to exactly two point attractors: a survival state, with AKT1 high, and an apoptotic state, with TP53 high and AKT1 low. Which one a patient reaches depends on the initial state, not on the input snapshot alone, so the switch carries hysteresis. That hysteresis is the object of control in Section 2.5, which works on the semi-tensor-product form of these rules, in which one synchronous update of a block is a single structure matrix acting on the lifted block

state.

One flag, *U*, records the uncoupled resistant configuration as a single bit per patient. It is read from the AKT and FOXO3a state that clinical work links to resistance [32, 33]. When *U* = 1 it holds AKT1 on and FOXO3a nuclear. The direction of information matters. *U* is read from the patient and encoded, not discovered by the model. Because *U* both derives from and acts on these nodes, the switch does not independently establish that a patient is resistant. For the same reason, the agreement in Section 3.2 between the survival attractor and the uncoupled state is a consistency check, not an independent validation, because both are computed from AKT and FOXO3a. What it adds is whether survival-axis inhibition moves a patient out of the coupled fraction, the quantity Section 3.4 reports.

### 2.5 Localised stabilisation and minimal-kernel design

The control objective is stabilisation to the apoptotic attractor. Global stabilisation would drive the network there from any initial state. On the full model this is not usable. The semi-tensor-product form lifts an *n*-node network to a 2^*n*^-dimensional map, and a delay of depth *d* raises that to 2^*nd*^ [37]. For 141 nodes with three timescales the lift is astronomical. We do not attempt it. We stabilise locally instead. A target set is locally stabilised when every trajectory from a neighbourhood of it converges to it under the free dynamics.

Local stabilisation becomes a network-level result through a decomposition. Partition the network into subsystems. Under weak coupling between subsystems, stabilising each subsystem locally, in sequence, drives the whole network to a neighbourhood of the joint target. This is the Lyapunov-based weak-coupling decomposition theorem of the BBCN118 framework [1]. Its formal statement and proof are restated in Appendix B. The design problem then has two parts. Choose a partition whose blocks are weakly coupled. Then stabilise each block on its own. Everything turns on the first choice.

The obvious partition is by pathway module, and it is the one BBCN118 used. It fails here. The apoptosis core does not sit in a weakly coupled module. Section 3.1 shows it sits inside a dominant strongly-connected core: 16 of the 22 pathway modules are mutually coupled. The coupling between these modules is strong and bidirectional. The weak-coupling premise of the theorem is violated for the pathway-module partition. Pathway-wise stabilisation therefore does not compose on the hard case. This is the obstruction of Section 3.1, seen from the control side.

We change the partition. Instead of splitting by pathway, we split by process timescale. The nine switch nodes fall into three blocks: fast (AKT1, PTEN, MTOR, PDPK1, PIK3CA; phosphorylation), mid (MDM2; turnover), and slow (TP53, PHLPP, FOXO3; transcription). The multirate schedule updates the fast block often and the slow block rarely, in the ratio 1:5:25, and the slow regulators enter the fast rules at a delay. This is what makes the blocks weakly coupled. Between two slow updates the slow block is constant. Over that interval the fast block evolves under fixed slow inputs, so it sees the slow block as a frozen parameter, not as a co-evolving variable. When the slow block does update, it reads a fast block that has already settled. The coupling does not vanish. It acts across timescales rather than within one. At any single timescale each block is weakly coupled to the others. This is the discrete analogue of timescale separation in singular perturbation. We therefore take the weak-coupling premise to hold block by block, where it failed on the pathway partition. We do not prove the coupling bound for this network. The singular-perturbation reading motivates it, and, as set out below, the endpoint on the full network is checked directly rather than assumed.

We therefore stabilise the switch block locally to its apoptotic attractor. The kernel is the set of clamped nodes that achieves this. We find it with the algebraic forward-stabilisation test [2]. From the block’s logical transition matrix *L* in semi-tensor-product form and a candidate clamp set *S*, form the state-modifying matrix 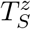 that pins *S* to its target values, the controlled matrix 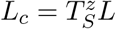, its transient power *ρ*, and the stabilisability matrix 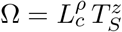. The set *S* stabilises the block if and only if every column of Ω is the target index vector. That is, clamping *S* drives every block state to target, not only the patient’s current state. Candidate sets are searched in increasing size. The smallest stabilising set is taken. Ties among minimal sets are broken by least causal score, then lexicographically, which is a tiebreak only, since every such set already stabilises. The search is summarised step by step in Appendix C.

The tractability follows from the partition. The test runs on a nine-node block, not on the full network. Its lift is 2^9^, small enough that the delayed algebraic form stays within reach where a 141-node lift would not. This is the dimension explosion of delayed semi-tensor-product design [37], avoided by confining the design to the switch. The clamp set the test returns is minimal and druggable. It is about two to three nodes, concentrated on the PI3K/AKT/mTOR axis.

Two points fix the scope of the claim. First, the kernel clamps only druggable nodes. It does not hold the downstream executioner nodes on by force. Once the switch is driven to the apoptotic state, commitment is reached through the intrinsic caspase cascade under the free dynamics, and the apoptotic state is a genuine fixed point, not a forced one. Second, the decomposition theorem certifies convergence to a neighbourhood of the target under weak coupling. It does not by itself certify the endpoint on the full network. We verify that endpoint directly. Section 3. shows that the switch-designed kernel commits the full 141-node network to the apoptotic fixed point, certified on the free cascade, patient by patient across the three cohorts. The design is local. The verification is global.

### 2.6 Scoring death: two readouts and the durability test

Death is scored two ways, and we report both. The commitment readout scores death by caspase commitment: CASP3 = 1. The strict readout adds that survival signalling is off: CASP3 = 1 and AKT1 = 0. The strict readout is the more conservative of the two.

A patient is durable when the free dynamics of the full 141-node cascade settle to a fixed point at which CASP3 = 1. The fixed-point test is one synchronous update of the free cascade that leaves the state unchanged. No node is held on. The apoptotic state is reached because PUMA, NOXA, and SMAC repress the survival guardians through the rules, and the executioner loop CASP6 → CASP8 → CASP3 self-sustains. It is not reached by forcing any executioner node. The durable state is therefore a genuine fixed point of the free dynamics, not a forced one. Because no node changes under the update, it is a fixed point under any update order, synchronous or asynchronous alike, so the durability verdict does not depend on the update scheme.

Durability is always certified on the free cascade. Every durable state reported here is a fixed point of the free cascade, reached under the rules, with nothing held on.

### 2.7 Reproducibility

Every number, figure, and table regenerates from the public repository with one command. The runs are deterministic, so the same inputs always give the same numbers. The pipeline self-verifies against a committed reference, so a drift in any figure is caught rather than hidden. The manuscript reads its numbers from the generated files, not from hand-typed values, so the text and the code cannot disagree.

The source-data terms are respected. Only the TCGA-BRCA binarised inputs are shipped with the code, since TCGA is open. TCGA alone therefore runs end to end out of the box. METABRIC is under controlled access, through the European Genome-phenome Archive, with processed data via cBioPortal under its data-use terms. I-SPY2 expression is available from GEO series GSE194040. For those two cohorts the repository documents the binarisation step, the robust *z*-score of Section 2.1, rather than shipping the patient-level matrices. An approved user regenerates byte-identical inputs and reproduces every downstream number. For the controlled-access cohorts, the repository also ships the derived-results files from which the reported numbers regenerate, so number-generation reproduces even without the patient-level data.

## 3 Results

### 3.1 Phenotypes are individually reachable but not jointly

The whole-network controller reveals the obstruction. Each cell-fate phenotype is reachable on its own. Driving apoptosis to a genuine fixed point succeeds in 86% of TCGA, 85% of METABRIC, and 81% of I-SPY2 patients. Proliferation-off succeeds in 94%, 95%, and 94%. The rates sit close together across three platforms, so individual reachability is a property of the network, not of one cohort. Resistance-off is the hardest phenotype everywhere. That is the first sign that the survival axis is held in place by the rest of the network.

The picture inverts when the three are pursued together. Each new phenotype must be reached without giving up the ones already won. Joint success collapses to 3% in TCGA, 2% in METABRIC, and 3% in I-SPY2 (Table 2). The gap between the high isolated rates and the low joint rate is the obstruction. Each phenotype is feasible. They cannot be held at once. Success is a genuine free-dynamics fixed point with no node forced, so this is not an artefact of pinning too few nodes. It is a statement about where the network can rest.

**Table 2.** Setup A controllability (% of patients reaching a genuine fixed point).

| Cohort / regime | Resistance-off | Apoptosis-on | Proliferation-off | All three |
| --- | --- | --- | --- | --- |
| TCGA, isolated | 8 | 86 | 94 | n/a |
| TCGA, sequenced | 8 | 9 | 3 | 3 |
| METABRIC, isolated | 4 | 85 | 95 | n/a |
| METABRIC, sequenced | 4 | 5 | 2 | 2 |
| I-SPY2, isolated | 8 | 81 | 94 | n/a |
| I-SPY2, sequenced | 8 | 9 | 4 | 3 |

The reason is structural. We build a directed graph on the 22 pathways, with an edge from one to another when a node of the first appears in a rule of the second, and compute its strongly-connected components [9]. One component dominates: 16 pathways that all feed back into one another. About 82 percent of pathways lie inside cyclic cores rather than a feed-forward chain (Figure 2). The coupling is distributed, not funnelled through one hub. Removing the strongest carrier, AKT1, shrinks the dominant component only from 16 pathways to 14. No one or two ablations break the cycle. This is the regime where pathway-wise stabilisation fails. Inside a strongly-connected core the other members re-inject the signals an intervention removes, so a phenotype can be feasible yet unreachable under any bounded budget. This is the weak-coupling premise of the decomposition theorem failing on the pathway partition (Section 2.5). It is why the apoptosis core must be modelled on its own terms.

**Figure 1.**
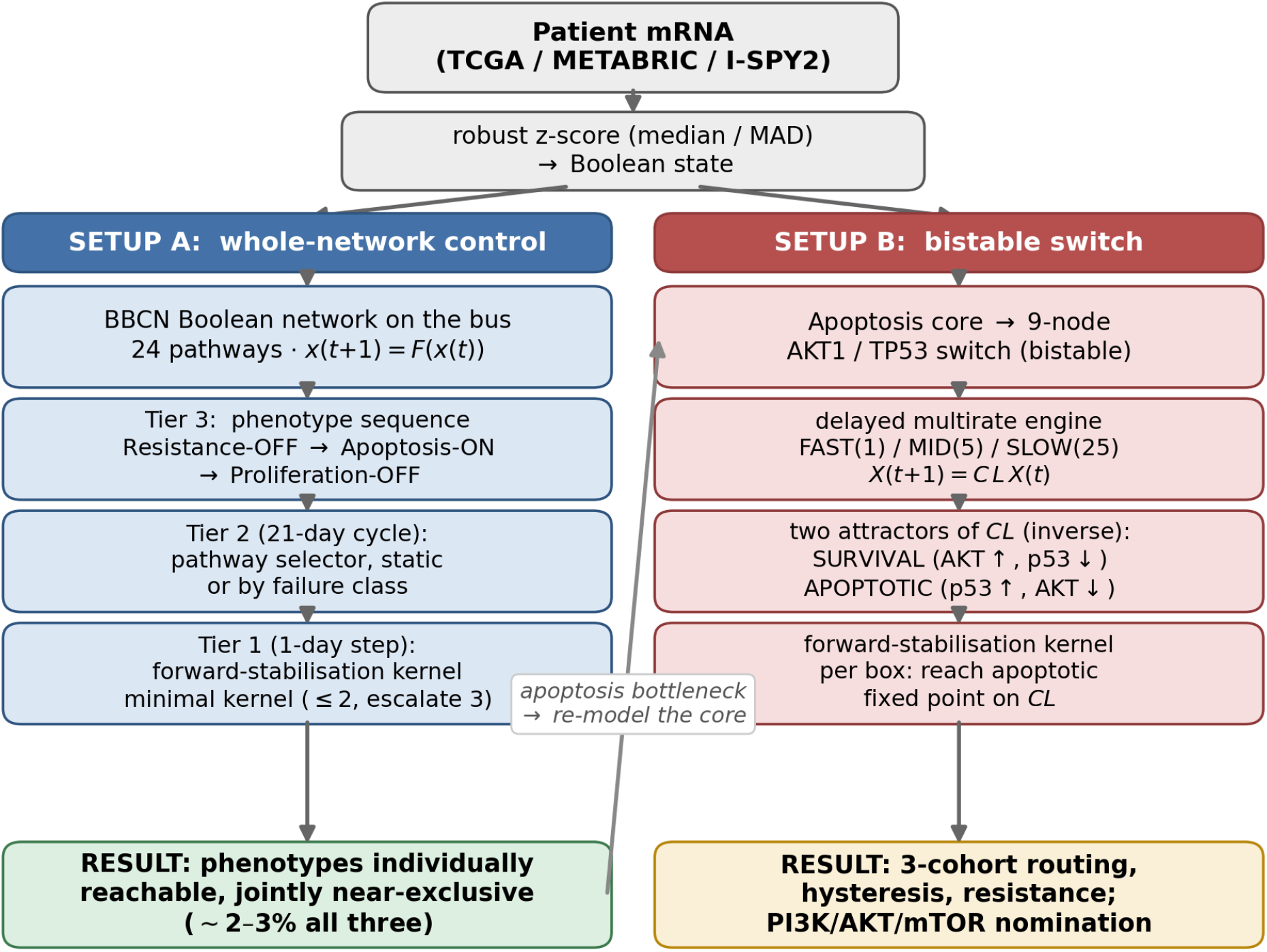
The BBCN control-design pipeline. A shared front end binarises patient mRNA into the initial state (Section 2.1). The whole-network controller (left) drives the full 141-node network under a capped, no-forcing budget and reveals the obstruction: the cell-fate phenotypes are individually reachable but jointly near-exclusive, held in a dominant strongly-connected core (Section 3.1). This motivates re-modelling the apoptosis core as a nine-node AKT1 and TP53 bistable switch, expressed as a multirate Boolean system (fast/mid/slow on a 1:5:25 schedule) (Section 2.4). Localised stabilisation of the switch, with a minimal druggable kernel, then commits the full network to a durable apoptotic fixed point (Section 2.5).

**Figure 2.**
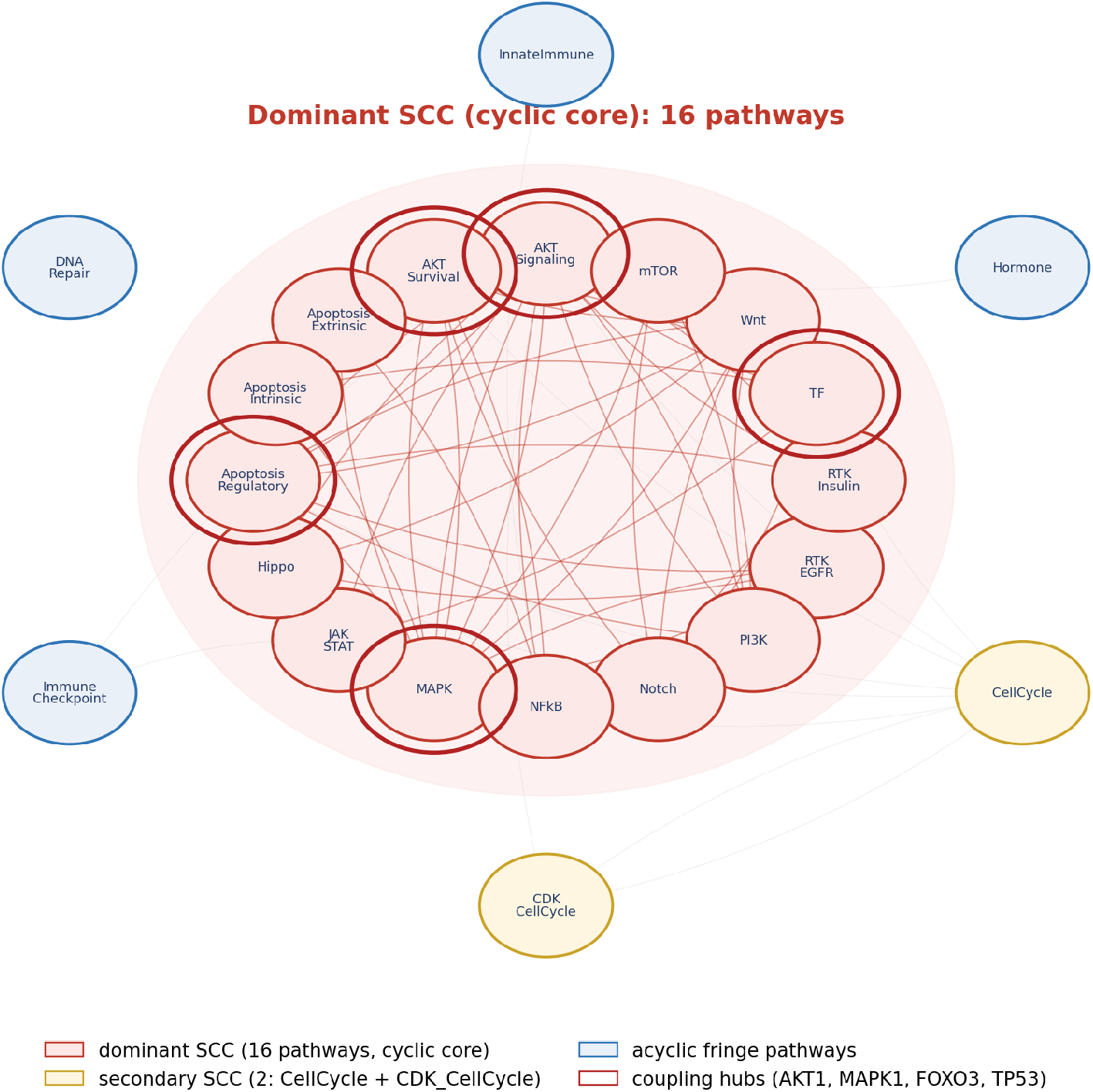
The pathway coupling graph. The dominant 16-pathway strongly-connected component is shaded as the cyclic core; the coupling hubs (AKT1, MAPK1, FOXO3, TP53) are ringed. The two auxiliary input modules are excluded.

### 3.2 The switch routes patients coherently across cohorts

Re-modelled as a bistable switch, the apoptosis core has two attractors. One is survival, with AKT high and p53 low. One is apoptotic, with p53 high and AKT low. The control problem becomes flipping the switch. Each patient’s binarised state seeds the switch, which runs on the multirate schedule until it settles. The switch routes about half of patients to the apoptotic attractor (50%, 48%, 51% for TCGA, METABRIC, and I-SPY2), about a quarter to survival, and the rest to a mixed outcome (Figure 3, Table 3). The split is stable across platforms.

**Table 3.** Setup B switch routing and biological check (robust *z*-score, full cohorts).

| Cohort ($N$) | Apoptotic | Survival | Mixed | SURV uncoupled |
| --- | --- | --- | --- | --- |
| TCGA (1082) | 50% | 36% | 14% | 95% vs 37% |
| METABRIC (1980) | 48% | 29% | 22% | 90% vs 29% |
| I-SPY2 (988) | 51% | 25% | 23% | 89% vs 26% |

**Figure 3.**
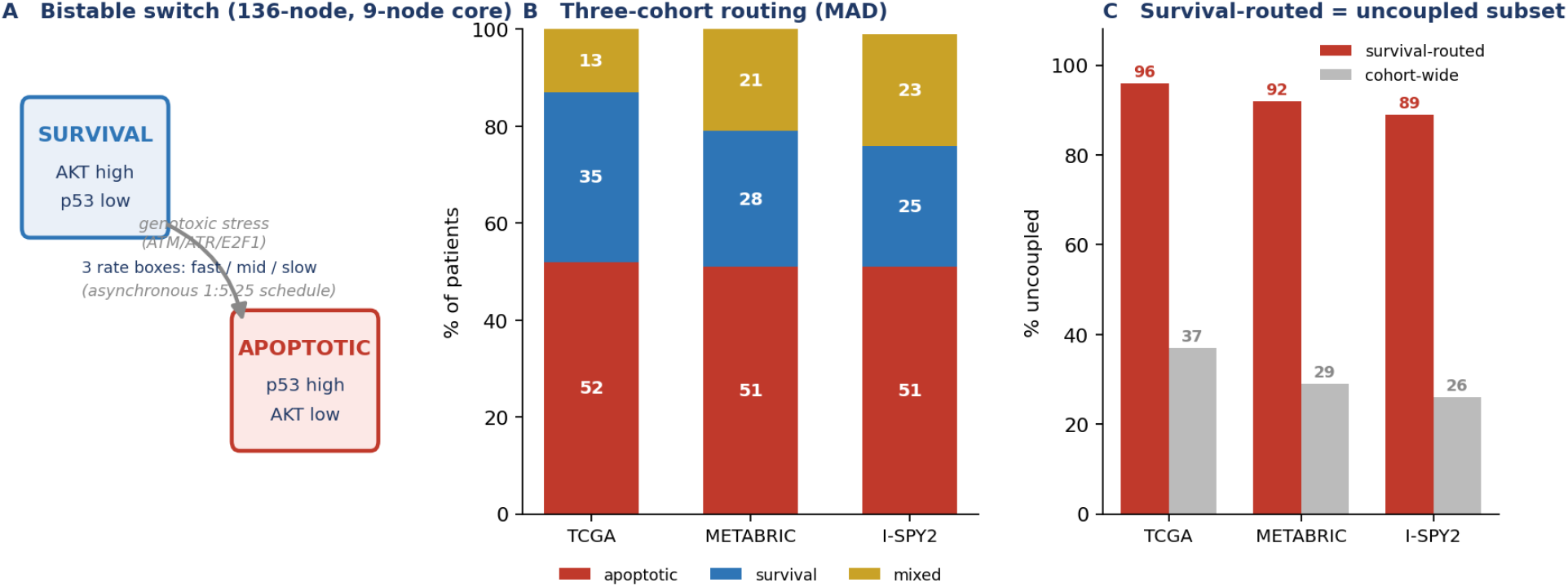
The bistable switch (141-node model). (A) The two attractors and the survival→ apoptosis flip under sustained genotoxic stress. (B) Three-cohort routing (robust *z*-score): about half apoptotic, a quarter survival, the rest mixed, consistent across platforms. (C) Biological check: survival-routed patients are 89 to 95% uncoupled, versus 26 to 37% cohort-wide.

The routing tracks biology. Survival-routed patients are 95%, 90%, and 89% in the uncoupled AKT and FOXO3a state, against cohort-wide rates of 37%, 29%, and 26%, an enrichment of about threefold. The switch sends the biologically resistant patients to the survival attractor. This agreement is expected, because the routing and the flag are both computed from AKT and FOXO3a. It is a consistency check, not an independent validation. A genuine test would use resistance markers from outside the switch.

### 3.3 Resistance is a hysteresis property with a coupling-state target

Survival-routed patients are not rescued by survival-axis inhibition on the baseline switch. Inhibiting the survival inputs does not flip the resistant attractor. Only a genotoxic input flips it. This is hysteresis. Once the uncoupled survival state is established, withdrawing the upstream drive does not cross back. Each timescale block is monostable on its own. The bistability, and this resistance, is emergent from the coupling between blocks.

The uncoupled state is not a discovery of the model. We encode it as the flag *U*, one bit per patient, from the binarised AKT and FOXO3a state, the uncoupling documented as a resistance mechanism [32, 33]. What the model contributes is to make *U* a control target. On the biology-faithful network, survival-axis inhibition durably ips the coupled fraction (33–36%), while genotoxic input is the weak lever (3–4%). Resistance corresponds to the uncoupled fraction, and the control target is the coupling state itself.

### 3.4 Localised stabilisation drives apoptosis on the full network

The kernel designed on the switch, by localised stabilisation, commits the full network to apoptosis. A minimal druggable kernel is found for 99–100% of resistant patients and concentrates on the PI3K/AKT/mTOR axis. Applied to the full 141-node network, the switch-designed clamp commits 92–95% of resistant patients to apoptosis, scored by caspase commitment. The design is local, on the nine-node switch. The effect is global, on the full network.

The network now settles into durable apoptotic fixed points. Re-running the staged controller on the biology-faithful branch, the full network reaches a durable apoptotic fixed point in 15–17% of patients, where the baseline supported none (Table 4). Every durable state is certified on the free cascade, as a genuine fixed point with nothing forced (Section 2.6).

**Table 4.** The biology-faithful network, all three cohorts. Drug-resistance equals AKT-FOXO3a uncoupling for 99.98% of 4050 patients. A designed PI3K/AKT/mTOR kernel commits 92–95% of resistant patients to apoptosis (commitment readout). The staged controller recovers durable apoptotic fixed points (15–17%) absent from the baseline network. All values are macro-sourced from `numbers2.tex` resistance, kernel, and durability are full-*N* runs, with the staged-controller figures re-verified on a per-cohort sample. Resistant, Durable P, Apoptosis-on, and Proliferation-off are percentages of the full cohort; Kernel commit is a percentage of the resistant patients.

| Cohort | Resistant | Kernel commit | Durable FP | Apoptosis-on | Proliferation-off |
| --- | --- | --- | --- | --- | --- |
| TCGA | 48% | 93% | 17% | 78% | 86% |
| METABRIC | 49% | 92% | 15% | 76% | 87% |
| I-SPY2 | 52% | 95% | 16% | 72% | 84% |
| Baseline | — | — | 0% | 81–86% | 94–95% |

We score death two ways and report both. The strict readout requires the executioner caspase on and the survival kinase stably off, and reads 8–14%. The commitment readout requires sustained caspase-3 activation, the point of no return, and reads 92–95%. The two differ because in uncoupled cells the survival kinase can reactivate after withdrawal, so the strict readout records a survivor even though the cell has committed. A committed cell does not un-die when a kinase switches back on. Both readouts stand far above the baseline network’s zero.

### 3.5 Minimality: a switch kernel matches near whole-network control

The durable state is secured by a far smaller intervention than the whole-network controller needs. We run both designs through the same delayed engine and the same release-and-persist test, scoring durability as persistence of death after every kernel node is unpinned. Death is reached in 100% of patients under either design. What separates them is whether it persists. The switch kernel holds the apoptotic state about as durably as the near whole-network controller (93.0/89.4/91.8% versus 94.0/93.4/93.9%). It is an order of magnitude smaller, about 2.6 versus about 31.99 nodes (Figure 4, Table 5). This is the paper’s minimality result. The switch kernel holds the same durable state with a focused, druggable intervention, where the whole-network controller needs near whole-network control that is not a deliverable regimen.

**Table 5.** Durability and kernel size, switch versus whole-network controller (release-and-persist test on the switch-routed resistant set). Source: cde_vs_switch_summary.csv.

| Quantity | TCGA | METABRIC | I-SPY2 |
| --- | --- | --- | --- |
| Resistant (switch-routed) | 202 | 398 | 238 |
| Kerneled (design found) | 199 | 376 | 231 |
| Death reached (held) % | 100 | 100 | 100 |
| Switch durability % | 93.0 | 89.4 | 91.8 |
| Whole-network durability % | 94.0 | 93.4 | 93.9 |
| Switch kernel size (nodes) | 2.6 | 2.52 | 2.52 |
| Whole-network size (nodes) | 31.99 | 29.62 | 33.26 |

**Figure 4.**
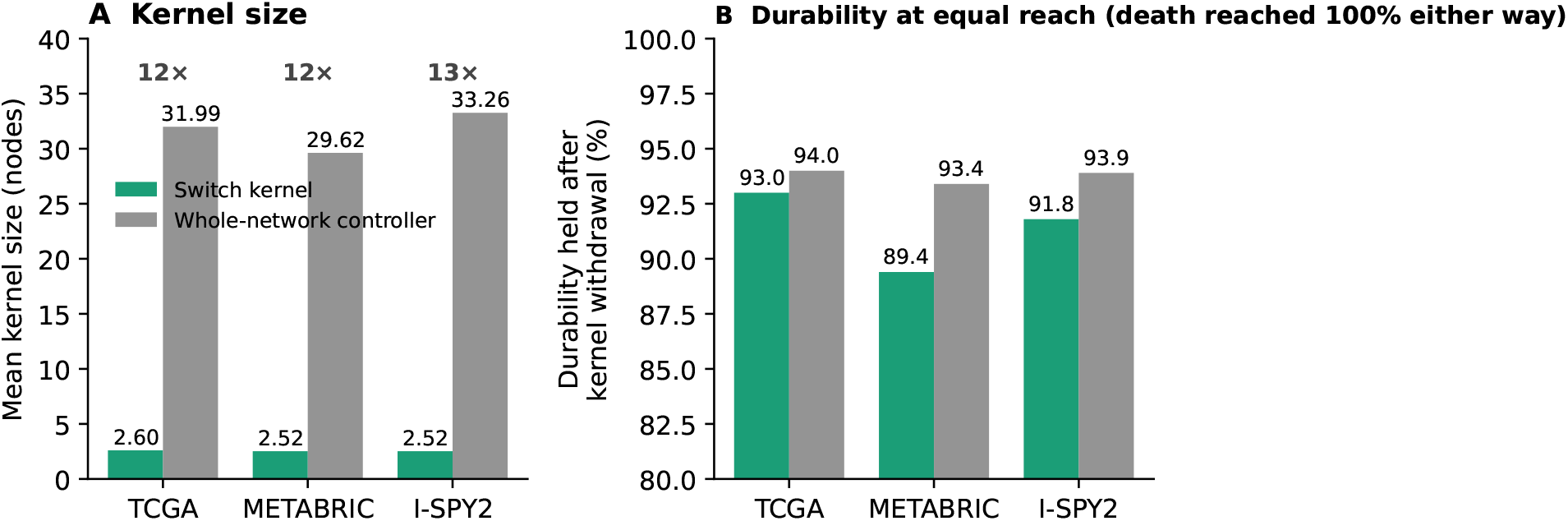
A minimal switch kernel secures the same durable apoptotic state as near whole-network control. (A) Mean kernel size, about 2.6 nodes for the switch versus about 31.99 for the whole-network controller, a roughly 12-fold reduction (12–13-fold across cohorts). (B) Durability after kernel withdrawal is essentially equal, with death reached in 100% of patients either way. Regenerated by `fig6.py` from `setup_a/data/cde_vs_switch_summary.csv`.

### 3.6 Drug-target nomination on the Pl3K/AKT/mTOR axis

The switch kernel concentrates on one axis. Tallying the switch-designed kernels by node, PIK3CA leads in every cohort, followed by AKT1 and the PDPK1 and MTOR nodes, with PTEN least used. The ordering is the same across all three cohorts and essentially identical under both kernel methods. The nomination is robust to the cohort and to the kernel-selection criterion. Separately, the whole-network controller yields a per-pathway nomination across the full pathway set, mapping each pathway’s kernel to a drug class and a lead agent (Table 6). That menu is broader than the switch kernel and spans pathways beyond the PI3K/AKT/mTOR axis. It is also robust to the data front-end. Repeated through median-binarised expression, multi-gene activity signatures, and phospho-protein activity, the PI3K/AKT/mTOR nomination holds.

**Table 6.**
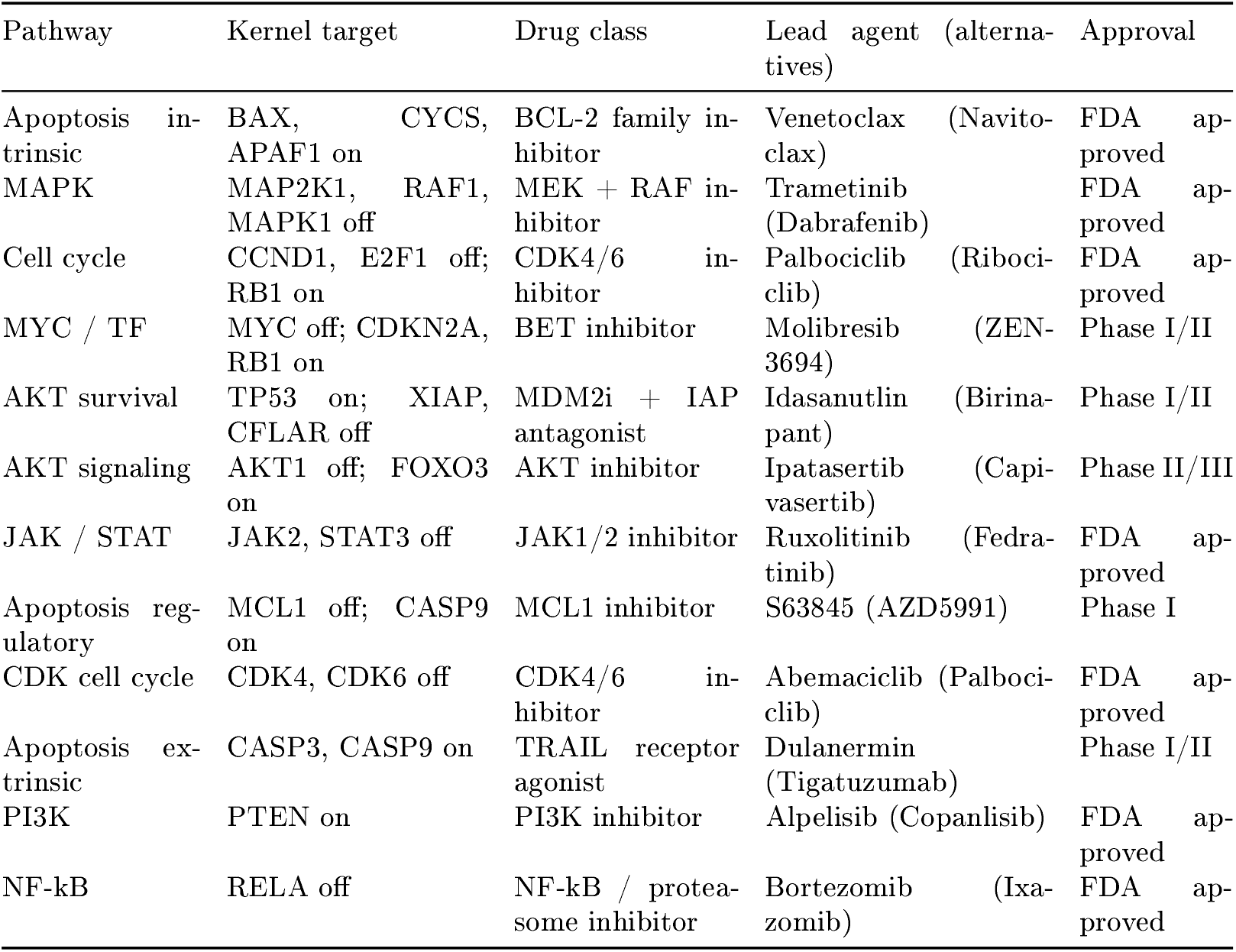
Kernel-to-drug nominations across the pathway set (source: `bbcn/drugs.py`). These mappings are illustrative structural nominations, one representative agent per pathway kernel they are not efficacy claims, and no drug was tested here.

The scope is stated plainly. The model is a structural drug-target nominator, not a response predictor. No survival benefit is claimed. The nominated axis was largely not administered to these chemo-era patients, so there is no response-validation cohort. The analysis is on primary tumours only.

## 4 Discussion

This paper makes one control-design claim, and it frames everything else. On a strongly coupled Boolean cancer network, global stabilisation is out of reach. Localised stabilisation is enough. Stabilising an extracted bistable switch to its apoptotic attractor drives the whole network’s phenotype, with a minimal druggable kernel.

The claim rests on a choice of decomposition. Our earlier weak-coupling decomposition theorem is exact under one condition: the coupling between subsystems must be small [1]. The natural subsystems are the pathway modules. On the apoptosis core that choice fails, because the core sits inside a dominant strongly-connected component and the modules are strongly coupled. We do not abandon the theorem. We change the partition. Splitting the network by process timescale, into fast, mid, and slow blocks, makes the coupling between blocks weak, because the delay embedding means each block sees the others as fixed over its own update interval. The theorem that failed on the pathway partition holds on the timescale partition, under a weak-coupling premise we argue but do not prove for this network. The transferable idea is that when a coupled network resists global control, the remedy may be a better decomposition rather than a bigger controller, and timescale separation is one that a delayed system supplies for free.

The first supporting result is why the switch is necessary at all. Cell-fate phenotypes in this network are individually reachable but jointly near-exclusive. They share a dominant strongly-connected core, so feasibility does not imply joint reachability. This complements stable-motif and semi-tensor-product control theory, which establish when a target attractor is controllable in principle. It adds that under a realistic capped budget on real patient states, the binding constraint is joint reachability across coupled phenotypes. That constraint is what forces the retreat to the core.

The second is that the hardest phenotype is best understood not as a marker to push but as one attractor of a bistable switch. Flipping it means crossing a hysteresis barrier that survival-axis inhibition alone cannot cross. Embedding biological timescale separation as a delayed multirate structure is what makes the core controllable where the whole-network controller could not. It also keeps the design tractable. The semi-tensor-product form of a nine-node block is small, where the lift of the full network, worsened by delay, is not. Confining the algebra to the switch is what avoids the dimension explosion of delayed Boolean control [37].

The third result concerns the network, and it is a statement about biology, not about our own model. A durable apoptotic fixed point exists only when the p53 feedback, the AKT and FOXO3a loop, and the commitment machinery are all in place biologically. With a degenerate p53 response, an unclosed AKT and FOXO arm, and no commitment step, no durable death state exists. With the p53 sensor, the loop closed by PHLPP, and the intrinsic caspase cascade in place, it does, and no parameter is tuned to make it appear. Durability is therefore governed by feedback fidelity, not by intervention strength. Drug resistance then corresponds to a precise molecular state, AKT and FOXO3a uncoupling, an established mechanism [32, 33] that the model makes a control target rather than a description.

A natural objection is that durability does not matter if a drug is never withdrawn. It does matter. Continuous inhibition is itself what breeds resistance, because sustained blockade relieves the pathway’s own feedback and the target reactivates [25]. A state held only by constant force collapses at every missed dose. A durable state inside the death basin survives them. Whole-network control across about thirty nodes is lifelong polypharmacy and is not tolerable, so minimality is what makes any maintenance conceivable. And apoptosis is terminal, so a cell that durably commits needs no further dosing. In this model that durable commitment is reached for a minority of patients, the 15–17% with a durable apoptotic fixed point; the argument here is about the value of durability where it is achieved, not a claim that it is achieved for most. The bistable-caspase basis of this commitment picture is developed by Eissing and colleagues, Legewie and colleagues, and Bagci and colleagues [27, 28, 29], with single-cell dynamics quantified by Albeck and colleagues [31]. The control-kernel and feedback-vertex-set view is due to Kim, Park, and Cho and to Fiedler and colleagues [30, 24], and the delayed Boolean control formalism to Li and Sun and to Cheng and colleagues [34, 35, 36].

The scope is stated plainly. The model’s outputs are a patient stratification, which is consistency-checked rather than independently validated, because the coupling ag is read from the same AKT and FOXO3a state the switch routes on, so an independent test needs resistance markers measured outside the switch; and a PI3K/AKT/mTOR drug-target nomination that is stable across the input front-ends we tested. The model does not predict pathologic complete response, which we attribute to endpoint distance rather than signal quality. It is a mechanistic hypothesis generator and a structural stratifier. Clinical benefit is not demonstrated here. The uncoupling flag is a proxy read from expression, invasion is out of scope, and the analysis is on primary tumours only.

One limitation is foundational, and we state it plainly. The model is single-cell and bistable, but the cohorts are bulk mRNA, each value an average over millions of cells. Near a bistable boundary an average need not correspond to any real cell: a tumour that is part survival-state and part apoptotic-state averages to an intermediate expression that names neither attractor. Binarising that average to seed the switch is therefore a coarse-graining, and it fixes the claim ceiling. The model is a patient-level stratifier and a structural target nominator, quantities a population average can support, not a per-cell attractor predictor, which it cannot. We have seen this directly. In bulk mRNA the coupling ag does not separate survival, and the phospho-AKT to nuclear-FOXO3a association the mechanism requires is close to zero at the population level. Both are expected if averaging collapses the switch, and both indicate where validation must happen: the coupling state should be read per cell, by co-localisation of phospho-AKT and nuclear FOXO3a on the same tissue section, not inferred from a bulk correlation that averaging washes out.

Two steps would raise the translational ceiling. The first is a durability-matched external test. Because durability corresponds to relapse avoidance, the direct test is whether AKT and FOXO3a coupling status, read from expression, stratifies relapse-free or overall survival in a long-follow-up cohort. That test should be run against the present model and reported at face value, whichever way it lands. The second is a fully cascade-native controller, in which the kernel search runs on the free cascade rather than on the engine used here for search only. It recovers additional fixed points and is the natural next development of the design.

## Author contributions

A.I.B. is the sole author and is responsible for the study design, the analysis, the software, and the writing of the manuscript.

## Code and data availability

All numbers, figures, and tables regenerate from the companion repository (https://github.com/AameriqbalBhatti/bbcn_public_repo) with a single command (`python run_all.py; python run_all.py --quick` for a fast check), which auto-detects the cohorts present. TC A-BRCA is open access and its binarised inputs are shipped with the code, so the pipeline runs out of the box on that cohort. METABRIC is available under controlled access through the European Genome-phenome Archive, with processed data via cBioPortal under its data-use terms. I-SPY2 expression is available from Gene Expression Omnibus series GSE194040. To comply with those terms the patient-level matrices for these two cohorts are not redistributed, but the deterministic binarisation is specified in Section 2.1, so a user with the primary data regenerates the inputs and reproduces every result.

## Declaration of generative AI use

During the preparation of this manuscript the author used a large language model (Anthropic’s Claude) as an assistive tool for L^A^TEX formatting and typesetting, for writing figure-generation code, for organising the companion code repository, and for language editing and drafting assistance on author-directed text. The tool was not used to generate scientific content, data, analyses, results, or their interpretation, and it is not an author. The author directed all such use, verified every reported number against the source code, and takes full responsibility for the entire content of the manuscript.

## A The BBCN 141-node model

This appendix lists the complete Boolean update logic of the 141-node BBCN model, one block per pathway module. For each node, the right-hand side is its update rule over intra-pathway nodes and external bus inputs; `AND`, `OR, NOT` are the Boolean connectives, and a node whose rule equals a single input is a pass-through of that input. Two held, per-patient inputs enter the rules: `U`, the uncoupled-state flag, set to 1 when the patient’s AKT activity and nuclear FOXO3a co-occur; and `GENOTOXIC`, the DNA-damage treatment input, off by default.

~~~
RTK EGFR. (Upstream_RTK)
EGF = EGF
EGFR = (EGF OR (ERBB2 AND ERBB3)) AND NOT MAP2K1
GRB2 = IRS1 OR (LATS1 OR (EGFR OR SRC))
GRB7 = GRB7 AND NOT PRKACA
ERBB2 = ERBB3 OR EGF
ERBB3 = NRG1 OR ERBB2
RTK INSULIN. (Upstream_RTK)
INS = INS
INSR = INS OR FOXO3
IRS1 = NOT RPS6KB1 AND (INSR OR IGF1R) PIK3CA = GRB2 OR IRS1
ABL1 = GRB2 OR EGFR OR ERBB2 OR IRS1
HORMONE. (Upstream_Hormone)
AR = FOXO3 OR NOT AKT1 PGR = ESR1
ESR1 = (FOXO3 OR KMT2D) AND NOT AKT1
KMT2D = PAX7
PI3K. (Terminal_support)
PTEN = NOT SRC
PDPK1 = (PIK3CA AND NOT PTEN) OR IGF1R
RAC1 = (PIK3CA OR DVL1) AND NOT AKT1
NOS3 = HSP90AA1 OR AKT1
MAPK. (Resistance)
HRAS = SOS1 AND NOT NF1 MAP2K1 = RAF1
RAF1 = HRAS OR KRAS KRAS = SOS1
SOS1 = GRB2
NF1 = (TP53 OR FOXO3) AND NOT (MAPK1 OR AKT1)
MAPK8 = (FAS OR TRADD) AND NOT (AKT1 OR MAPK1)
MAPK1 = MAP2K1 AND NOT DUSP1
DUSP1 = MAPK1 OR MAPK8 OR JUN
WNT. (Invasion_OFF)
APC = AXIN1
AXIN1 = NOT DVL1 AND GSK3B
CTNNB1 = NOT (APC AND AXIN1 AND GSK3B) AND (WNT1 OR SRC OR YAP1)
DVL1 = FZD3 OR NF1
FZD3 = WNT1 OR CREB1
RND1 = GRB7 OR DVL1
TCF7L2 = CTNNB1 OR JUN
WNT1 = WNT1
NOTCH. (Invasion_OFF)
ABL2 = ABL1
DLL1 = DLL1
HES1 = NOTCH1
HEY1 = NOTCH1
EIF4EBP1 = NOT MTOR
MYOD1 = NOT (HES1 OR ABL1)
MYOG = NOT HEY1 AND MYOD1
NOTCH1 = DLL1 OR CTNNB1 OR LCK OR MDM2
JAK STAT. (Xglobal_Tier1)
JAK2 = IL6 AND IL6R
STAT3 = JAK2
STAT1 = JAK2 AND NOT AKT1
SOCS3 = STAT3 OR STAT1
HIPPO. (Invasion_OFF)
YAP1 = NOT LATS1
LATS1 = NF2 OR LCK
NF2 = NEDD4L
AKT SIGNALING. (Xglobal_Tier1)
AKT1 = ((MTOR AND PDPK1 AND PIK3CA AND NOT PTEN) AND NOT PHLPP) OR (U AND AKT1)
FOXO1 = NOT (SGK1 OR AKT1)
SGK1 = MTOR OR PDPK1
NEDD4L = NOT (SGK1 OR PRKACA)
FOXO3 = (NOT AKT1) OR U
PHLPP = FOXO3 AND NOT U
GSK3B = NOT (AKT1 OR MAPK1)
MTOR. (Invasion_OFF)
RHEB = NOT TSC2
MTOR = RHEB OR (PIK3CA AND NOT PTEN)
TSC2 = NOT (AKT1 OR MAPK1) AND (PRKAA2 OR GSK3B)
TWIST1 = MAPK1 OR AKT1 OR MAPK8
TF. (Xglobal_Tier1)
CEBPB = ABL2 OR CREB1
CREB1 = AKT1 AND NOT PTEN
EIF4E = NOT EIF4EBP1
GATA3 = NOT STAT1
CXCL8 = NOT GATA3
MYC = (CTNNB1 OR PIM1 OR PIM2 OR PIM3 OR MAPK1 OR STAT3 OR ESR1) AND NOT GSK3B
NFKB. (Resistance_OFF)
IKBKB = TNF OR IL1A OR IL1B OR TLR2 OR TLR4 OR AKT1
NFKBIA = NOT IKBKB
RELA = IKBKB AND NOT NFKBIA
INNATE IMMUNE. (Inflammatory)
IL1A = IL1A
IL1B = IL1B
TLR2 = TLR2
TLR4 = TLR4
AKT SURVIVAL. (Apoptosis_ON)
MDM2 = (TP53 OR AKT1) AND NOT (CDKN2A AND (E2F1 OR ATM OR ATR OR GENOTOXIC))
TP53 = (NOT MDM2) OR ATM OR ATR OR GENOTOXIC
JUN = (MAPK8 OR MAPK1 OR RELA) AND NOT (PPARG OR DUSP1)
XIAP = (AKT1 OR JUN) AND NOT SMAC
CFLAR = (AKT1 OR RELA OR CREB1) AND TP53
APOPTOSIS REGULATORY. (Apoptosis_ON)
MCL1 = (NOT GSK3B AND MAPK1) AND NOT NOXA
PAK1 = TCF7L2
PRKACA = AKT1 OR WNT1
SRC = PRKACA OR ABL1
APOPTOSIS EXTRINSIC. (Xglobal_Tier1)
FAS = FASLG
FASLG = JUN OR RELA OR TP53
FADD = FAS OR TRADD
TRADD = FAS
CASP8 = (FADD AND NOT CFLAR) OR CASP6
BID = CASP8
CASP3 = (CASP8 OR CASP9) AND NOT XIAP
CASP9 = CYCS AND APAF1
CASP7 = (CASP8 OR CASP9) AND NOT XIAP
CASP6 = CASP3 OR CASP7
APOPTOSIS INTRINSIC. (Apoptosis_ON)
BAD = NOT (PAK1 OR AKT1 OR MAPK1 OR PIM1 OR PIM2 OR PIM3)
BAK1 = (BCL2L11 OR TP53 OR BCL2) AND NOT (MCL1 OR BCL2 OR BCL2L1)
BAX = (BCL2L11 OR TP53 OR BCL2) AND NOT (MCL1 OR BCL2 OR BCL2L1)
BCL2 = (NOT (BAD OR BCL2L11) AND (CREB1 OR MAPK1)) AND NOT PUMA
BCL2L1 = (NOT (BAD OR BCL2L11) AND CREB1) AND NOT PUMA
BCL2L11 = FOXO3 OR FOXO1
CYCS = BAK1 OR BAX
APAF1 = TP53 OR FOXO1 OR FOXO3
SMAC = BAX OR BAK1
PUMA = TP53
NOXA = TP53
CELLCYCLE. (Proliferation_OFF)
CDKN2A = NOT (MYC OR SRC OR AKT1 OR MAPK1)
CCND1 = (TCF7L2 OR TEAD1 OR MYC) AND NOT (CDKN2A OR CDKN1A OR GSK3B)
E2F1 = NOT RB1
CDKN1A = TP53 OR FOXO3 OR NOT (MAPK1 OR AKT1 OR MYC)
RB1 = NOT (CDK4 OR CDK6 OR CDK2 OR CCND1)
TEAD1 = YAP1
CDK CELLCYCLE. (Proliferation_OFF)
CDK4 = CCND1 AND CDKN1B
CDK6 = CCND1 AND CDKN1B
CDK2 = CDK4 OR CDK6
CCNE1 = CDK4 OR CDK6
CDKN1B = FOXO1 OR FOXO3
DNA REPAIR. (DNA_Repair)
ATM = ATM
ATR = ATR
BRCA1 = ATM OR ATR
BRCA2 = BRCA1
PARP1 = PARP1
CHEK1 = ATM
CHEK2 = ATR
RAD51 = BRCA1
IMMUNE CHECKPOINT. (Immune_Checkpoint)
PDCD1 = LCK
CD274 = IFNG OR STAT3 OR AKT1
AUX INPUTS1. (Boundary)
HSP90AA1 = HSP90AA1 (external input)
IGF1R = IGF1R (external input)
IL6 = IL6 (external input)
IL6R = IL6R (external input)
LCK = LCK (external input)
NRG1 = NRG1 (external input)
PAX7 = PAX7 (external input)
IFNG = IFNG (external input)
AUX INPUTS2. (Boundary)
PIM1 = PIM1 (external input)
PIM2 = PIM2 (external input)
PIM3 = PIM3 (external input)
PPARG = PPARG (external input)
PRKAA2 = PRKAA2 (external input)
RPS6KB1 = RPS6KB1 (external input)
TNF = TNF (external input)
~~~

*Total: 141 nodes across 24 modules, of which 22 are regulatory pathways and two are boundary blocks carrying external inputs. The pathway graph of Section 3*.*1 is built on the 22 regulatory pathways*.

## B The weak-coupling decomposition theorem (prior work)

This appendix states the weak-coupling decomposition theorem on which the localised-stabilisation design of Section 2.5 rests. The theorem and its proof are the contribution of the earlier BBCN118 preprint [1] (CC BY 4.0) the statement and proof sketch below are reproduced and adapted from that work, so that Section 2.5 can be read without consulting the source. The remark at the end is the new content. It records that the theorem fails on the pathway partition but holds on the timescale partition, which is the move the paper makes.

**Setup**. Let a Boolean network with global update *x*(*t*+1) = *F* (*x*(*t*)) be partitioned into *m* pathway modules *P*_1_, …, *P*_*m*_, with module states *x*_*i*_ and module targets 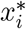, so that *x* = (*x*_1_, …, *x*_*m*_) and 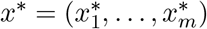.

Theorem l (convergence under pathway decomposition)

Assume:

(A1) *Weak inter-pathway coupling*: 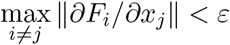, with *ε* ≪ 1

(A2) *Local kernel effectiveness:* for each module *i* there is a kernel *u*_*i*_ that drives 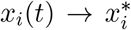, in finitely many steps and

(A3) *Modularity*: *F*_*i*_(*x*) ≈ *f*_*i*_(*x*_*i*_, *e*_*i*_(*x*)), where *e*_*i*_(*x*) collects the inputs *P*_*i*_ receives from other modules.

Then the sequential application of the pathway kernels 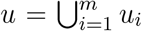 drives the global state into a neighbourhood of the target,

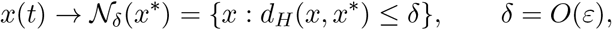

where *d*_*H*_ is the Hamming distance.

Proof sketch

Take the weighted Hamming Lyapunov function

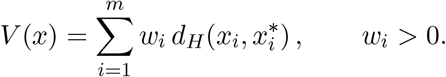

*V* is non-negative, integer-valued, bounded above by ∑_*i*_ *w*_*i*_|*P*_*i*_|, and zero exactly at *x*^∗^. Intervening on module *k* reduces its term by some *α*_*k*_ *>* 0 by (A2) . By (A1) every other module moves by at most *w*_*j*_*ε*|*P*_*j*_|, so

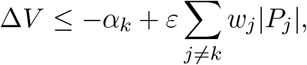

which is strictly negative once *ε* is small enough relative to *α*_*k*_. Because *V* is bounded below, strictly decreasing, and integer-valued, it reaches a minimum *V*_min_ in finitely many steps, with *V*_min_ ≤ *δ* = *O*(*ε*) at *ε* = 0 the bound gives *V*_min_ = 0 and exact convergence. □

### Corollary l

(convergence rate). If each pathway kernel reduces the mismatch by at least *β >* 0 per intervention, the system reaches the neighbourhood in at most *T* ≤ *V* (*x*(0)) */* (*β/*2) interventions.

### Remark

the partition is what matters. The result holds only under assumption (A1) . On the pathway partition the present network violates it. Its pathway-coupling graph carries a dominant strongly-connected component of 16 of the 22 regulatory pathways (Section 3.1), so the inter-module coupling is not small and pathway-wise stabilisation does not deliver joint reachability. The paper does not abandon the theorem. It re-partitions the apoptosis core by process timescale, into fast, mid, and slow blocks. The delay embedding makes each block quasi-static over the others’ update intervals, so the inter-block coupling satisfies (A1) where the inter-pathway coupling did not, and the theorem applies block by block (Section 2.5) . The full proof, including the explicit *ε* threshold, is in the cited preprint and is not reproduced here.

## C The forward-stabilisation kernel search

Figure 5 sets out the kernel search of Section 2.5 step by step, in the form in which it is implemented.

**Figure 5:**
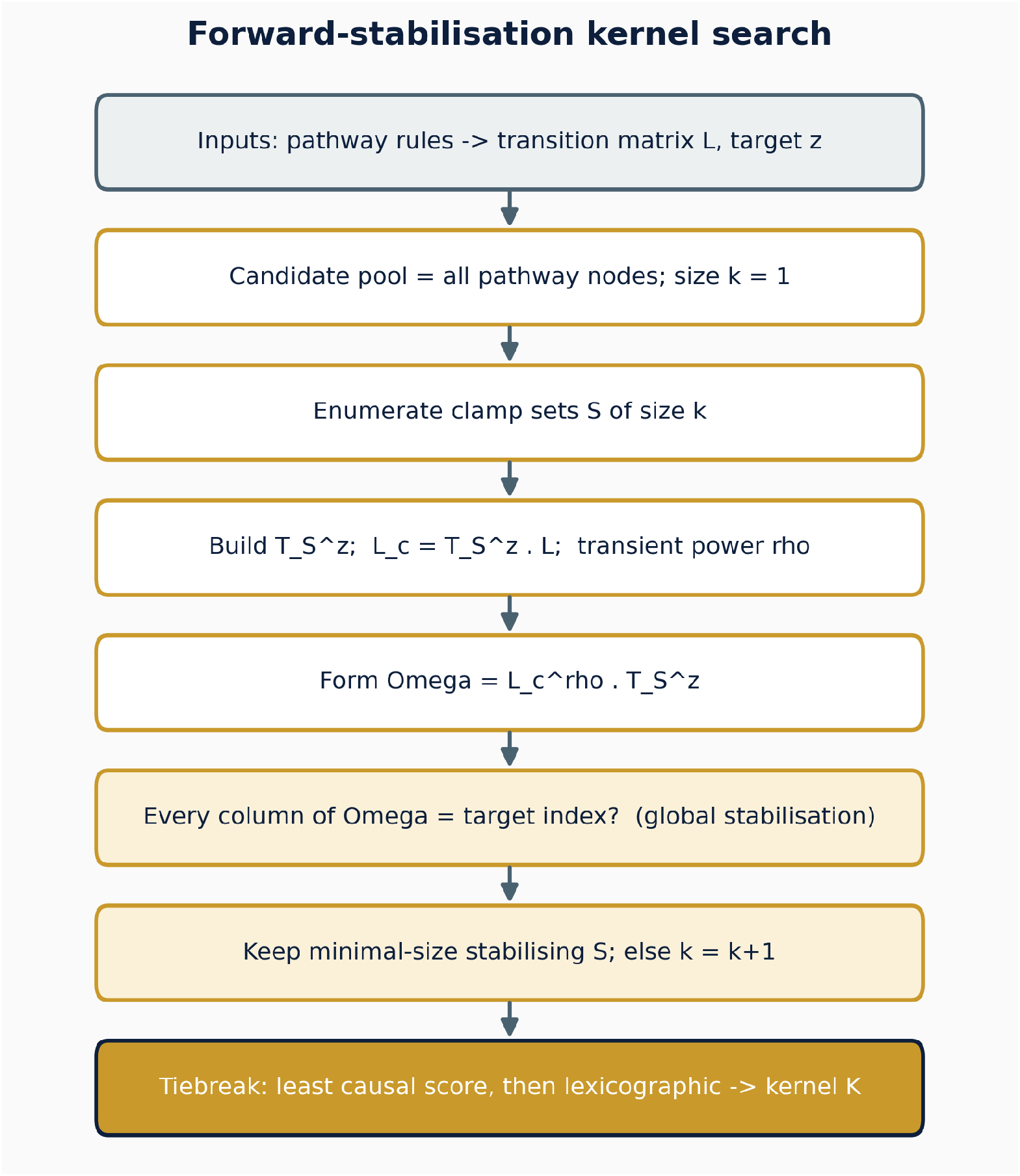
The forward-stabilisation kernel search. Candidate clamp sets are enumerated in increasing size. or each set the algorithm forms the clamped operator and accepts the set only if it drives every state of the block to the target, not only the patient’s current state. The smallest accepted set is kept and ties are broken deterministically. The test is tractable on the nine-node switch block, whose lift is 2^9^, and not on the full 141-node network.

