## Supplemental Table and Figure for "Localised stabilisation of a bistable switch for minimal-kernel phenotype control in a Boolean breast-cancer network" (from another version of this preprint: version 2, 2026-07-08)

### Supplementary Information
### The Boolean Breast-Cancer Network (BBCN), Paper 3 v2

#### S1. The two kernel methods and the two controllers, compared

This supplement gives the full comparison summarised in Section 3.8 of the main text. Both kernel designs, the ranked reachability heuristic and the algebraic global-stabilisation test (Methods, Tier 1), were run on both the static controller (which fixes the per-phenotype pathway set) and the dynamic whole-network controller (which selects pathways by failure classification), on all three cohorts at full $N$, on the bare, un-repaired 135-node network under synchronous updating. Table 1 reports the per-phenotype results; routing in Setup B is unaffected by the kernel method, since kernels are designed only after a patient is routed. In the figures the algebraic method is labelled *stabilize* and the heuristic *ranked*; these are two names for the same kernel-selection step.

Three readings follow. First, the ranked method reproduces the earlier BBCN118 kernels exactly, so the heuristic results are preserved rather than discarded. Second, on the static controller the two methods are close (apoptosis differs by at most a few points), because for the per-pathway targets used here the patient's own state already lies in the stabilising basin. Third, on the dynamic controller the algebraic method certifies fewer passes than the heuristic, the gap being largest on proliferation (TCGA 65% ranked versus 39% algebraic), because it demands that the clamp drive *every* pathway state to target, not only the patient's starting state. The algebraic figure is the conservative, provable one; the difference between the two columns is exactly the set of patients the heuristic counts from a favourable initial state that are not globally stabilisable. In Setup B the two methods give identical flip rates in every cohort (TCGA 47% vs 47%, METABRIC 42% vs 42%, I-SPY2 46% vs 46%), with PIK3CA the leading actuator throughout: on the small bistable boxes the algebraic test confirms the heuristic kernel rather than contradicting it.

Table 1: **Table S1.** Kernel-method and controller comparison (% reaching target), full cohorts, bare 135-node network. Static = fixed pathway set; Dynamic = whole-network failure-classification selection. Each cell pair is ranked | algebraic.

| | | Static controller | | Dynamic (whole-network) | |
|---|---|---|---|---|---|
| Cohort | Phenotype | ranked | algebraic | ranked | algebraic |
| TCGA | Apoptosis | 86 | 91 | 31 | 27 |
| | Proliferation | 94 | 85 | 65 | 39 |
| METABRIC | Apoptosis | 85 | 89 | 33 | 29 |
| | Proliferation | 95 | 91 | 61 | 40 |
| I-SPY2 | Apoptosis | 81 | 87 | 39 | 35 |
| | Proliferation | 94 | 91 | 60 | 43 |

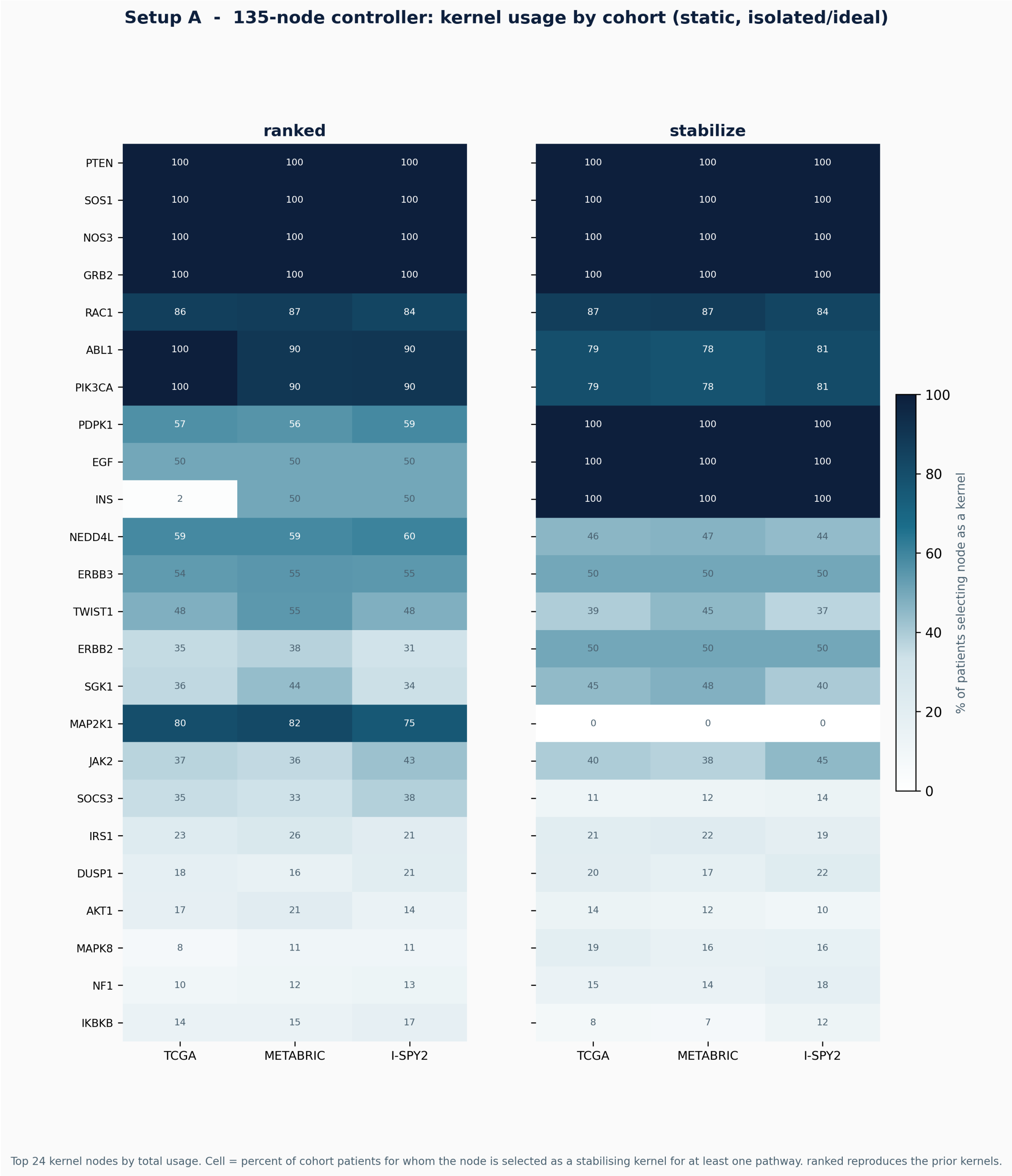


Figure 1: **Figure S1. Setup A kernel usage by cohort.** The 24 most-used kernel nodes under the static, isolated/ideal controller, shown for the ranked heuristic (left) and the algebraic (stabilize) test (right); each cell is the percentage of cohort patients for whom the node is selected as a stabilising kernel for at least one pathway. A small core, PTEN, SOS1, NOS3, and GRB2, is used for every patient in every cohort. The algebraic test sets MAP2K1 to zero and EGF and INS to full use, reflecting its stricter global-stabilisation criterion, while the dominant nodes are unchanged. The usage pattern is stable across TCGA, METABRIC, and I-SPY2.
